# HU searches and binds specific DNA via a multi-step process combining weak electrostatic binding, protein reorientation and DNA flexibility

**DOI:** 10.1101/2025.07.18.665543

**Authors:** Elliot W. Chan, Mark C. Leake, Agnes Noy

## Abstract

Despite their limited resolution, coarse-grained simulations have been the chosen method to obtain mechanistic information regarding the process of protein binding to DNA. Here, we demonstrate that state-of-the-art atomic simulations can capture binding events and provide new molecular insight. By using the bacterial model protein HU, we link existing crystal structures to the search and recognition states. We find that weak contacts mediated by only a few positively charged residues enable initial binding, followed by facilitated diffusion and transient rolling on DNA. Extended arms in the HU structure serve as antennae to search for DNA, permitting intrasegmental hops and intersegmental jumps. The transition to specific binding only occurs at the DNA target site, helped by its bendability, indicating a “concerted” binding mechanism between the protein and DNA. We observe a lack of direct binding to the most positively charged area, which is hidden behind HU’s arms and defines the recognition state. This prevents the protein from being trapped in random DNA. Instead, the multi-step binding process found here ensures that the high-affinity complex only occurs at the target position. We anticipate that other DNA-binding proteins with multiple surface-charged regions and DNA-bend induction capability might follow a similar strategy.

## Introduction

The mechanism by which proteins locate and attach to their specific binding sites on chromosomal DNA is crucial for virtually all genomic functions. In this process, the protein must find the target site within nonspecific DNA and establish binding through specific interactions [1]. The limitations in the resolution of experimental techniques in terms of time and space have made it difficult to characterize the mechanisms behind DNA-protein association [2, 3]. As a result, simulations have been employed to complement experimental studies, particularly using coarsegrained models due to their capacity to sample experimentally relevant timescales in contrast to all-atom simulations [4, 5, 6, 7].

The combination of the two approaches has proven valuable, revealing that the search for target sites is accelerated beyond 3D diffusion by a series of mechanisms categorized under the model of “facilitated diffusion” [8]. These include 1D sliding along DNA [9, 10], brief intrasegmental “hops” [11, 12], and intersegmental “jumps” between nearby DNA segments [3]. 1D sliding was characterized to follow the DNA grooves, hence being rotation-coupled due the structure of the double helix [2, 13, 14]. The initial assumption was that the protein would remain intimately associated to the grooves. However, the variation in binding energies across the different sequences for most of the proteins was quickly detected to be too large to allow for efficient searching [15, 16]. The presence of deep traps separated by large energy barriers is even more pronounced for architectural DNA-binding proteins as a result of the conformational alterations imposed on DNA [17].

To address the conflicting demands for speed and stability, a two-state model was proposed: a search state, characterized by weak interactions between the protein and DNA, allowing for efficient sliding; and a recognition state, where the protein forms a high-affinity complex with its specific DNA binding site [15, 10]. Consequently, rotation-coupled sliding occurs only when the two states are highly similar, albeit with weaker DNA:protein contacts in the searching one [2, 18, 19]. However, a large number of proteins, such as DNA architectural factors, possess numerous positively charged regions all over their surface that could serve as sliding interfaces [20, 21]. The existence of different areas for specific and non-specific binding could facilitate fast exploration, although it would introduce the necessity to transition between the various binding modes [10, 22]. Here, we employed atomic-precise simulations to elucidate the binding mechanisms for this prevalent class of proteins, which remain largely unknown. The incor-poration of greater detail on the molecules facilitated the characterisation of conformational transitions, as well as the role of various structural features, including the positively charged secondary regions and DNA deformability.

The DNA-binding protein HU was selected as a case study due to the abundance of single-molecule and structural data [23, 24, 17, 25]. HU is a highly abundant and conserved nucleoid-associated protein in bacteria, involved in numerous processes including transcription, replication, and repair [26]. It binds and bends DNA via its elongated *β*-ribbon arms, exhibiting a strong attraction for distorted structures such as nicks, kinks, and cruciforms (Figure 1A) [27, 23]. In addition, HU can bind B-DNA with modest affinity via the lateral side of its *α*-helix body, forming a complex that was embedded in extensive cooperative HU:DNA filaments as found in a crystal structure (Figure 1B) [24].

**Figure 1:**
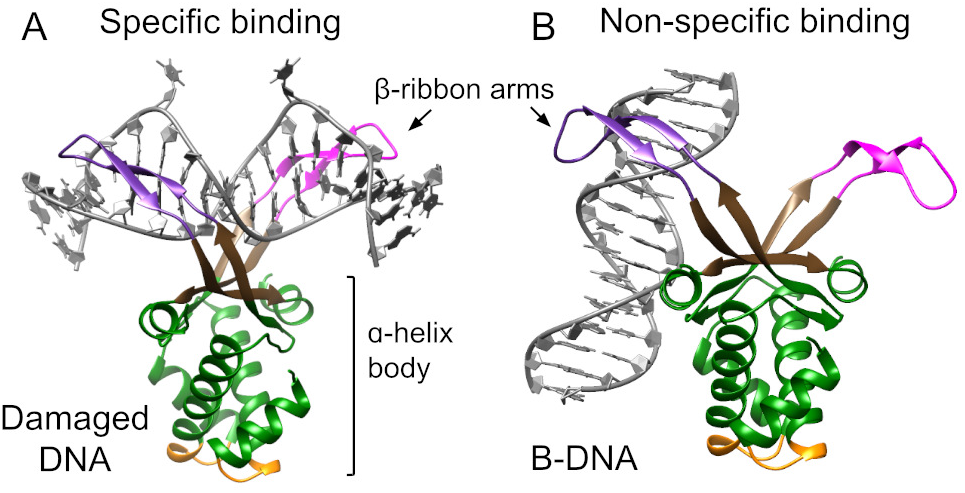
DNA binding modes as detected by crystallography and adapted to HU*αβ* heterodimer from *E. coli* (see methods for details): (A) specific binding to damaged DNA based on PDB 1P78 [23]; (B) non-specific binding to B-DNA based on PDB 4YEW [24]. Color marks HU regions: arm from HU*α* in magenta, arm from HU*β* in purple, saddle between arms in brown, protein body in green, and protein “bottom” in yellow.

Experiments have identified three diffusion rates for HU along DNA, associated with distinct sliding mechanisms: rotational-coupled for the slowest one, in which the DNA would be embraced by HU’s arms and the saddle between them; and hopping for the fastest one, which was thought to be mediated by the flexible arms [28, 17, 29]. Another study revealed that a series of lysines located on the HU’s body (Lys3, Lys18 and Lys83) were also important for 1D diffusion [25]. As a result, a rotational-uncoupled diffusion model was proposed, where the protein would slide using its lateral site, thus explaining the intermediate diffusion rate [25]. However, it is still unclear whether this lateral binding mode is adopted by a single HU unit.

Here, we demonstrate that state-of-the-art atomic simulations are now capable of providing new in-sights into the mechanisms behind DNA searching and recognition by proteins. Our simulations successfully captured numerous spontaneous association events between HU and DNA, including intrasegmental hops and intersegmental jumps. By reproducing the two experimental structures, our simulations replicated the formation of the non-specific complex, thus confirming its stability, as well as the transition to the specific one. We discovered that weak electrostatic interactions are essential to mediate the initial protein-DNA binding, which subsequently can progress to the non-specific complex due to the reorientation of the protein. We also observed that the transition to the specific complex only occurs in the presence of damaged DNA due to its increased flexibility, ensuring that this strong binding does not occur on random DNA. All these observations, derived from the atomic detail of our simulations, allow us to propose a structural model for the three diffusion modes of HU along DNA.

## Materials and Methods

The Amber20 software package was employed for configuring and executing simulations via the CUDA implementation of the pmemd program [30]. All simulations were solvated via TIP3P octahedral boxes (unless specified otherwise) with a minimum distance of 15 Å between the solute and the box’s edge. The systems were neutralized with a 0.2 M concentration of potassium and chloride ions determined by the Dang parameters [31]. The ff14SB and parmBSC1 forcefields were employed to represent the protein and DNA, respectively [32, 33]. Simulations were performed under constant T and P (300 K and 1 atm) following standard protocols [34, 35].

A complete structure of the HU heterodimer from *E. coli* was generated by fitting the protein’s *α*-helix body from PDB 4YEW [24] to the body from *Anabaena* (PDB 1P78) [23]. This allowed the incorporation of the flexible *β*-ribbon arms, which were unresolved in PDB 4YEW. The amino acids of the arms were then mutated to the *E. coli* sequence.

The DNA of PDB 1P78 was included to build the specific HU-DNA complex in which the protein interacts with damaged DNA (see Figure 1A). The sequence consists of CA**T**A*T*CAATTTG-T-TG, where *T* is unpaired and flipped, **T** is unpaired but stacked, T represents the central T:T mismatch and the – symbol indicates the position of the unpaired Ts in the complementary strand. The single-stranded nicks present in 1P78 were repaired to enhance the structural stability. The complete structure was then minimized and simulated for 5 ns, allowing us to evaluate the preservation of the *β*-ribbon secondary structure in the arms. This trajectory (designated as ‘expt-dmg’) served as reference for the experimental structure, as its short duration guaranteed that the complex remained in that state without undergoing a significant change.

The PDB 4YEW served as a framework to construct the non-specific HU-DNA complex. The coordinates of the complete *E. coli* HU heterodimer were superimposed to the protein’s body from 4YEW and the DNA was elongated at both ends to a total of 60 bp (see Figure 1B). The structure was then minimized and simulated for 5 ns, acting as reference for the experimental structure in which HU binds to B-DNA (expt-BDNA). The trajectory was then extended to 500 ns (this simulation is referred to as ‘BDNA’) to evaluate the stability of this conformational state. An equivalent structure was generated by incorporating the same damaged DNA as before at 8 bp off center to facilitate the transition from non-specific to specific binding. The simulation was extended to 2 *µ*s and designated as ‘damaged’.

The heterodimer HU was placed 3 nm away from a 60-bp DNA fragment with the same sequence as before (Figure 2A). This distance was selected to be sufficiently small to be covered during the simulation time, yet bigger than the Debye–Hückel screening length, allowing the protein some degree of translational and rotational freedom before interacting with DNA. The Debye–Hückel length is the distance at which two electric charges stop “sensing” each other due to the presence of other ions in solution. At the simulated salt concentration (200 mM KCl), this distance is ∼0.7 nm, which was less than a quarter of the initial separation between the protein and the DNA. The system was then simulated for 100 ns.

**Figure 2:**
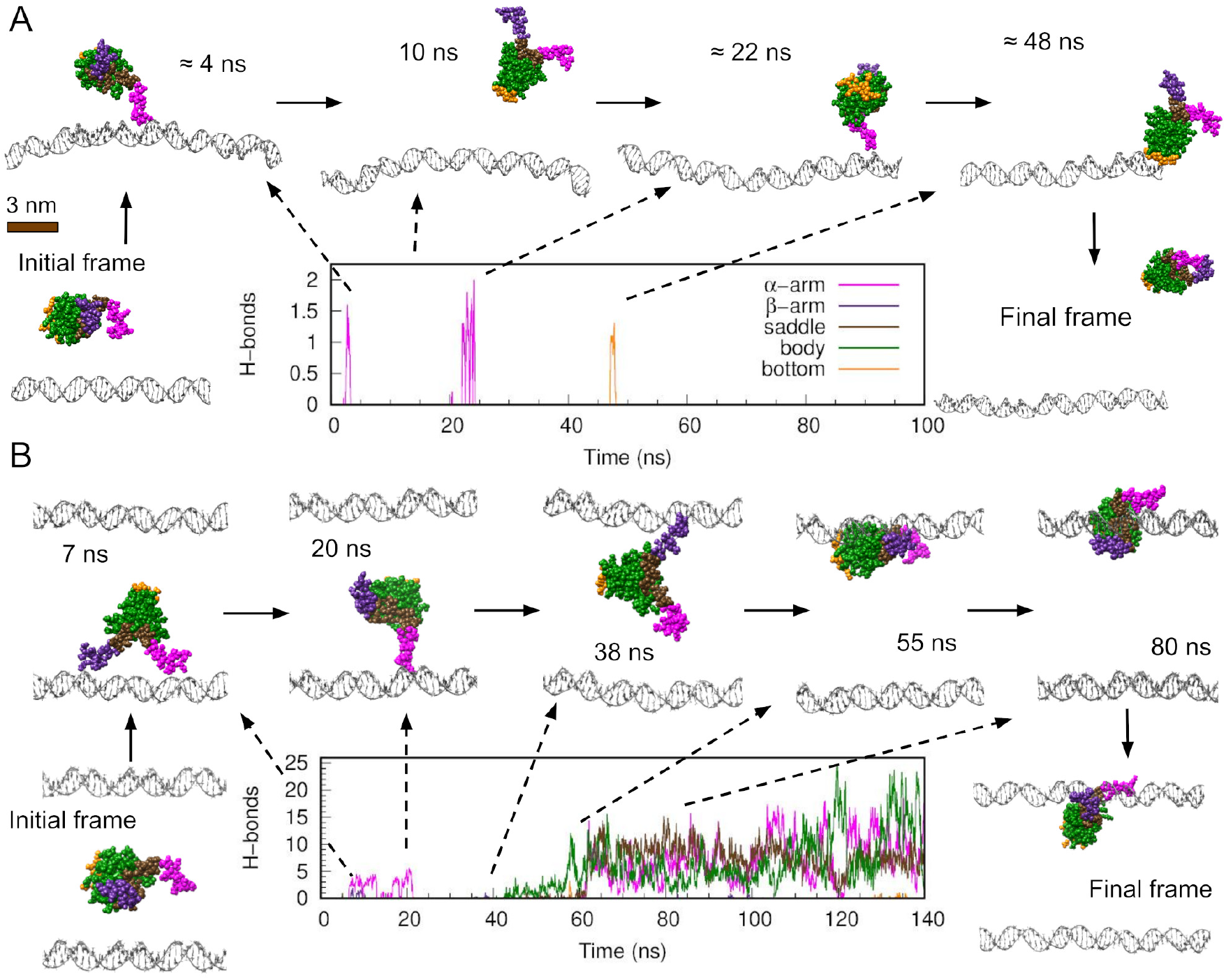
HU hops and jumps on DNA using its extended arms. Time evolution of hydrogen bond formation between DNA and various HU regions (color code as in Figure 1) together with representative frames from two independent simulations initiated with HU 3 nm away from DNA. (A) Preliminary simulation performed with a 60-bp DNA embedded in octahedral solvation box. (B) First replica of the ten performed with a 150-bp DNA solvated in a rectangular box, where the top duplex is the periodic boundary copy of the bottom one from the central solvation box.

After this preliminary simulation (called replica 0), we performed more replicas with an extended DNA fragment of 150 bp. To limit the size of the system, the DNA and HU (placed again 3 nm away) were solvated by a rectangular box rather than a truncated octahedron. The DNA ends were fixed to guarantee the molecule did not cross over into the next boundary box and produce large scale self-interactions. Ten independent replicas were initiated from this structure and were extended to 140 ns, based on the time at which HU and DNA reached a stable state that remained unchanged for 50 ns or the protein drifted away.

Hydrogen bonds were determined using the cpptraj program with a distance cutoff of 3.5 Å and an angle cutoff of 120 degrees [36]. The number of hydrogen bonds formed by each amino acid with DNA was capped to 1, hence the time-averages across the simulations reflect the ratio of frames exhibiting contact. We used our software, WrLINE/SerraLINE, to measure the DNA bending angles. First, we calculated the molecular axis for each frame with WrLINE [37], and then we extracted tangent vectors to calculate the bend angles with SerraLINE [38, 39]. The two tangent vectors were fitted using 16-bp fragments that were separated by 22 bp around the HU binding site. This approach was similar to the one previously used with PDB 1P78, which enabled to determine the overall bending induced by HU [23].

## Results

### HU hops and jumps DNA via its *β*-ribbon arms that serve as antennae

To determine whether simulations could capture binding events, we placed the protein HU 3 nm away from a 60-bp DNA fragment (Figure 2A). At the onset of this simulation (called replica 0), the protein quickly migrated towards the DNA, establishing a short-lived contact (lasting 2 ns) with one of its arms. The protein then diffused for approximately 20 ns before briefly interacting again via its arm, resulting in a hop along 20 bp away from the initial contact site (Figure 2A and movie S1). Afterwards, the protein detached, rotated, and briefly interacted with its “bottom” before drifting away. This third contact occurred at the same DNA site as the second one, which is likely due to the fact that the second interaction was already near the end of the DNA segment (Figure 2A).

After this preliminary simulation, we extended the DNA fragment to 150 bp in order to provide HU with additional space. In the first replica, HU made an intersegmental jump between copies of DNA that resulted from the implementation of periodic boundary conditions in our simulations (Figure 2B and movie S2). HU initially contacted the DNA from the central box before crossing the box’s boundary and interacting with DNA’s periodic copy. The proximity between DNA periodic copies (∼9 nm) resembles an environment of a high DNA density or long DNA molecules that are permitted to coil. Our simulation demonstrated that intersegmental jumps can occur relatively easily (in time scales of the order of ns) when two DNA fragments are in close proximity, which is consistent with previous experiments [3].

We observed that hopping and jumping were facil-itated by HU’s *β*-ribbon arms, in particular by Arg61 and Lys67 placed at the top (see Figure 3A). The elongated nature of HU’s arms enabled them to serve as antennae, detecting DNA fragments in the vicinity. The electrostatic attraction between negatively charged DNA and positively charged DNA-binding proteins generally promotes their initial approximation due to its long range. However, the distance at which they cease to ‘feel’ one another is less than 1 nm at physiological salt concentration, as determined by the Debye–Hückel screening length. With a length of around 3 nm, HU’s arms can explore areas beyond the reach of the protein’s body, roughly three times the Debye–Hückel screening length. Previous coarse-grained simulations detected that disordered protein tails facilitated intersegmental transfer thanks to a “monkey-bar” mechanism where the protein interacted with two DNA duplexes at the same time [40]. Here, we discovered that extended and flexible regions can also promote intersegmental jumps, with the protein temporarily detached, as they exert electrostatic attraction on DNA segments beyond the expected Debye limit.

**Figure 3:**
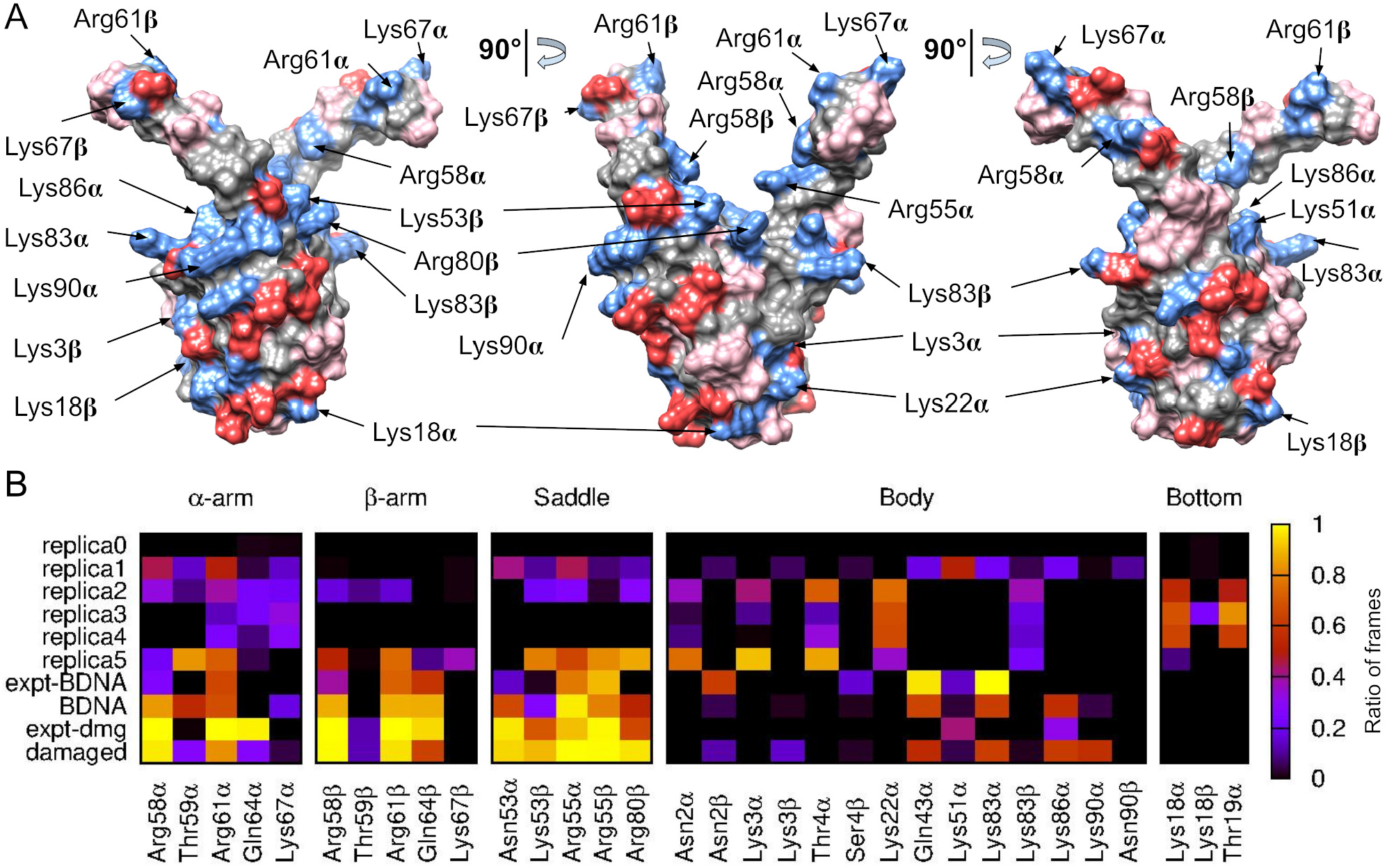
Structural details of *E. coli* HU*αβ* and its interaction with DNA. (A) Three views rotated by 90 degrees illustrating the physicochemical nature of HU’ surface residues: positively charged in blue, negatively charged in red, polar in pink, and apolar in grey. The aminoacids labelled are the positively charged ones making significant interactions with DNA. (B) Ratio of frames presenting hydrogen bonds between each HU’s residues and DNA for: the preliminary simulation started with HU positioned 3 nm from a 60-bp DNA (replica 0); five representative replicas of the ten started with HU 3 nm from a 150-bp DNA (replica 1-5); a short simulation (5 ns) started with HU bound to B-DNA as in PDB 4YEW (expt-BDNA); an extension of the previous simulation to 500 ns to examine potential transitions (BDNA); a short simulation (5 ns) started with HU bound to damaged DNA as in PDB 1PT8 (expt-dmg); and a very long simulation (2 *µ*s) started with HU bound to damaged DNA as in PDB 4YEW to examine transitions to the structure observed in 1PT8 (damaged). DNA-HU contacts in replica 0 are not visible in the heatmap due to their short duration.

### Weak initial contact leads to lateral binding via HU rolling over DNA

In replica 2, HU contacted DNA using one of the *β*-ribbon arms, as in the previous replicas (Figure 4A). This interaction served as an anchor point for subsequent interactions with the protein’s bottom and the rest of the body. HU then made a series of “rolling” events back and forward thanks to the transient dissociation of the arm’s contact, until the protein settled on the DNA through its lateral side (Figure 4A and movie S3). The final structures resembled the experimental one, where HU binds to DNA non-specifically (Figure 1B), forming more interactions with its *α*-helix body rather than with the saddle between the *β*-ribbon arms (Figure 4A) [24].

**Figure 4:**
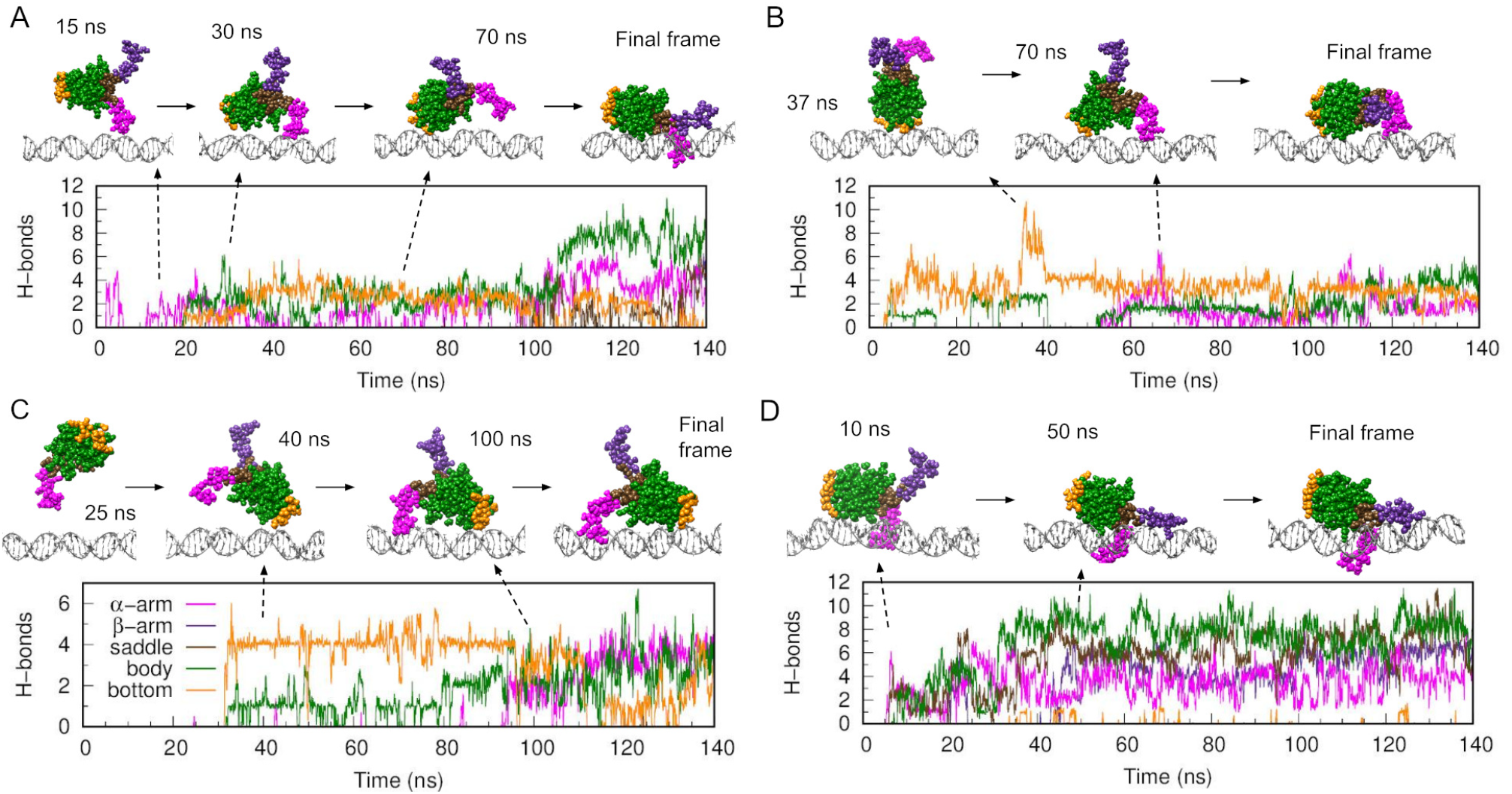
HU transitions from the encounter complex to lateral binding by rolling over DNA. (A-D) Time evolution of hydrogen bonds formation between DNA and various HU regions (color code as in Figure 1) together with representative frames from replicas 2 (A), 3 (B), 4 (C) and 5 (D) from the ten calculated using a 150-bp DNA solvated in a rectangular solvation box and initiated with HU 3 nm away. Replicas 6 and 7 are in Figure S1 due to their similar features, while replicas 8, 9, and 10 did not present contact between the protein and DNA.

The back-and-forth oscillations were also present in replica 6 (Figure S1) and could be the foundation of the “skipping” events seen in previous experiments [9, 22]. In these, the lac repressor was found to repeatedly traverse its target spanning 45 bp, before establishing the specific protein-DNA complex [9]. This length is consistent with our proposed model of the protein rolling over DNA, which would facilitate finding the most effective positioning between the two molecules for making strong interactions.

Replicas 3, 4, and 7 demonstrated that the encounter complex between HU and DNA could also be established by the protein’s bottom in addition to the *β*-ribbon arms (see Figure 3B, 3C and S1). These initial binding states remained stable for approximately 50 ns before transitioning to a lateral-binding conformation similar to the previous final state, where DNA interacts along the HU’s body (movie S4 and S5). The transition was made again by the protein rolling over DNA. The protein’s ability to reorient itself before making any contact with DNA also confirmed that the 3 nm starting distance was adequate to observe a variety of encounter complexes. For example, in replica 4, the protein rotated 180 degrees, positioning itself to the left instead of the originally selected right before interacting with the DNA. (Figure 3C and movie S5).

Finally, replica 5 confirmed the stability of the lateral binding pose, as the protein established direct contact with DNA in this state at around the 10th ns and maintained that position for over 100 ns (Figure 3D and movie S6). In replicas 8-10, the protein drifted away before any contact, and therefore, they were not included in our report.

### HU’s residues involved in the process of DNA binding

We then conducted a detailed analysis of the interactions between HU and DNA by identifying the amino acids responsible for establishing hydrogen bonds with the DNA (Figure 3B). To determine the interaction map of the conformations seen experimentally, we conducted two short additional simulations (lasting only 5 ns) of the structures derived from experiments where HU is already bound to either damaged or undamaged DNA, as shown in Figure 1 (designated as expt-dmg and expt-BDNA, see methods). While we found that expt-dmg uses the saddle area for interactions with DNA, expt-BDNA engages with both the saddle and the protein’s body, albeit predominantly with the protein’s body (Figure 3B).

In our replicas, interactions with the saddle were scarce due to its inaccessibility behind the protein’s arms. The exceptions were replicas 1 and 5, which presented the structures more similar to those of expt-BDNA. We saw that for most of our replicas, DNA interacted with the protein’s body in ways similar to expt-BDNA, depending on which side of the protein was touching the DNA. (see Figure 3B). These include residues already identified experimentally (Lys3 and Lys83), as well as others presented in both subunits (Asn2, Thr4) or exclusively in one subunit (Lys22*α*, Gln43*α*, Lys51*α*, Lys86*α*). Lys18 was the third residue experimentally identified for the lateral binding, and along with Thr19*α*, they contributed to stabilizing DNA interaction with the protein’s bottom (Figure 3B). Finally, due to their inherent flexibility, the two arms presented a diverse pattern of interaction across the different simulations.

### DNA bendability facilitates transition to specific binding

To further assess the stability of the lateral binding, we extended the expt-BDNA simulation to 500 ns. We observed that the protein temporarily shifted toward binding the DNA with the two arms and saddle in an attempt to transition to the specific binding mode; although, it quickly reverted to a configuration resembling that at the beginning of the simulation (Figure 5A and movie S7).

**Figure 5:**
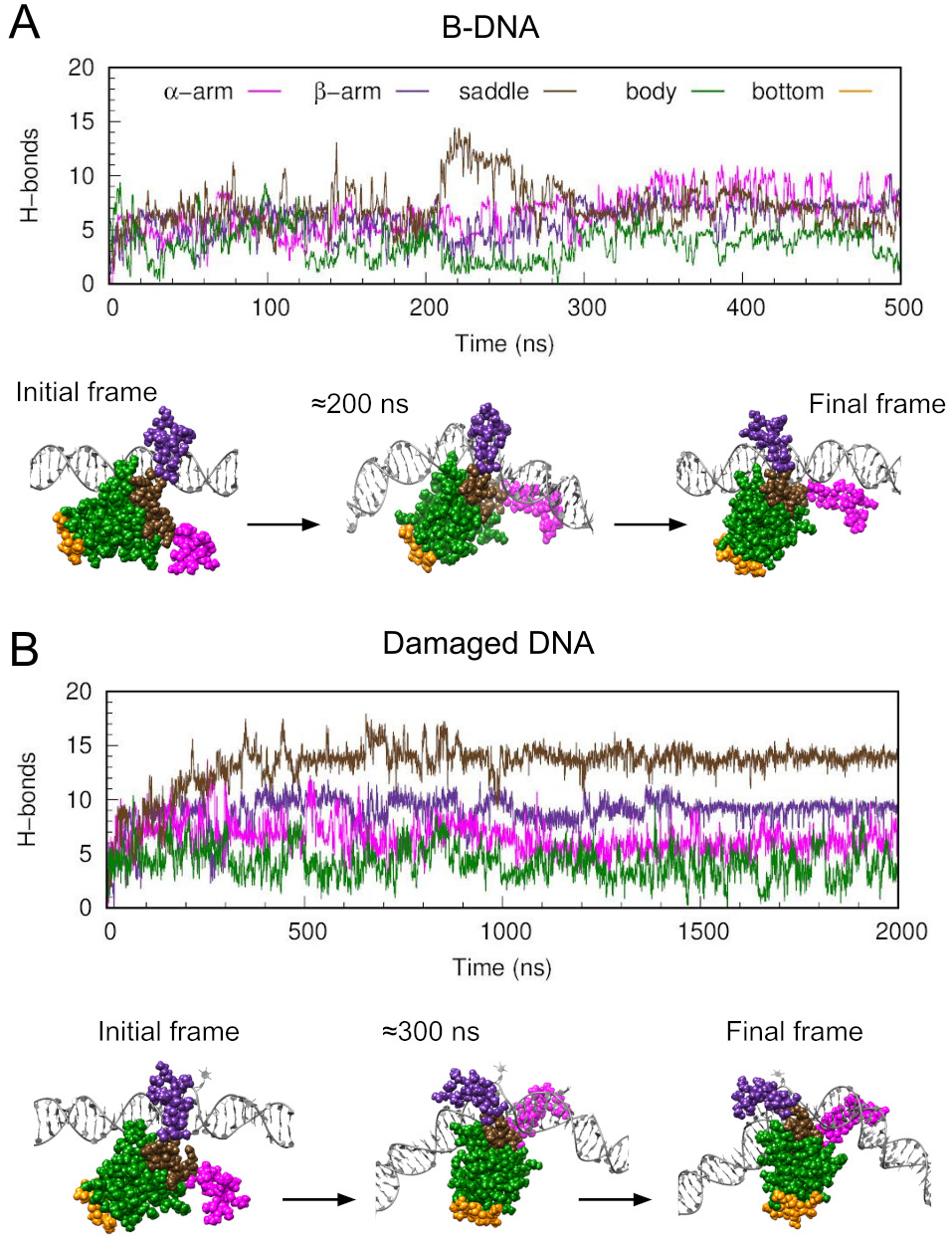
HU transitions from non-specific to specific binding in the presence of damaged DNA. Time evolution of hydrogen bond formation between DNA and various HU regions (color code as in Figure 1) together with representative frames for: (A) the simulation started with HU bound to B-DNA as in PDB 4YEW (designated as ‘BDNA’ in Figure 3B); and (B) the simulation started with HU bound to damaged DNA as in PDB 4YEW and changing to the structure observed in 1PT8 (designated as ‘damaged’ in Figure 3B).

We then introduced DNA damaged into the existing structure in a attempt to induce the transition to the specified binding mode. The defects were the same as in PDB 1P78 (see methods), and were located where they could be recognized if the shift was successful. In the initial 400 ns, HU entered to a mode reminiscent of the specific binding where the hydro-gen bonds with the saddle surpassed those with the protein’s body (Figure 5B). We extended the simulation to 2 *µ*s to allow to the second arm to move inside the DNA’s minor groove, thereby aligning with the specific binding mode; although it remained trapped interacting with the DNA backbone (movie S8). Nevertheless, the similarity in the HU:DNA contact maps between this simulation and expt-dmg indicated that the protein was already well positioned relative to the DNA, and it was just a matter of time to complete the transition (see Figure 3B).

DNA bending was evaluated for the two simulations and compared with two extra simulations containing the same DNA fragments (damaged and undamaged) but without HU. The bending of damaged DNA without HU was 34 ± 17 degrees (mean ± standard deviation), which was slightly higher than the bend of naked B-DNA (28 ± 13 degrees). These values increased to 55 ± 23 degrees for damaged DNA and to 34 ± 16 degrees for B-DNA in the presence of HU. Nevertheless, they were still far from the bend observed in the crystal structure (around 100 degrees). These findings indicate that, while HU’s arms are crucial for causing the strong DNA bending of specific recognition, some flexibility in the DNA itself facilitates making this bending possible. In the context of the “induced-fit” and “conformation selection” scenarios, we found a middle ground that we call the “concerted” mechanism, where the natural flexi-bility of damaged DNA helped kick off the change, allowing the protein to bend the DNA more easily.

## Discussion

Here, we demonstrated that all-atom simulations can provide novel, relevant insights into the process of target searching and specific binding (Figure 6). We looked at the HU protein, which can bind DNA in two forms as have been detected experimentally (Figure 1): to damaged DNA with high affinity, using its extended arms and the saddle between them [23]; and to the rest of the DNA with low affinity, using its *α*-helix body [24]. The non-specific binding was detected on a cooperative nucleoprotein filament, and so its existence on a single HU was an open question [24].

**Figure 6:**
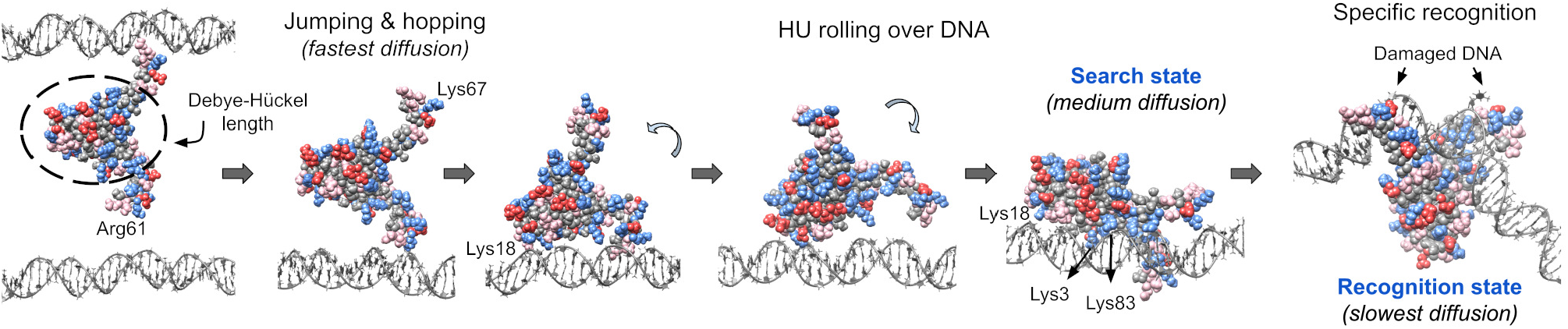
Model of the multi-step process of HU binding to DNA. The positively charged amino acids at the tip of the protein’s arms facilitate searching for DNA beyond the Debye-Hückel length set by its body (shown approximately by a dashed-line ellipse), allowing for hopping and jumping. Following initial binding, HU reorients itself over DNA (depicted by 3D arrows) until it reaches the search state, which is secured by a triad of lysines (3, 18, and 83) [25] among other residues. When damaged DNA is present, HU reaches the recognition state, in which DNA attaches to the most positively charged area (the saddle between arms). We predict that these three stages correspond to the three diffusion rates discovered in experiments [29]. Overall, HU and DNA bind in a multi-step process, where electrostatic interactions are progressively increased, ensuring the high-affinity complex only happens at the specific binding site.

We discovered that HU’s protruding arms functioned as antennae to sense nearby DNA beyond the Debye–Hückel limit of protein’s body, enhancing HU’s ability to hop and jump within DNA (Figure 6). The two positively charged residues (Arg61 and Lys67) at the top of the arms were the ones responsible for making these interactions, despite being surrounded by negative charges and nonpolar chains (see Figure 3A). Similarly, different parts of the protein, such as the bottom, could also form encounter complexes with a limited number of positive charges. We thus found that large positive patches are not required for establishing encounter complexes, and that they may even be detrimental due to slowing down the searching dynamics [41]. These first contacts were stable enough to hold the DNA, but weak enough to not trap it in this initial configuration (see Figure 6).

The encounter complexes subsequently evolved to the lateral binding via HU’s *α*-helix body, thus confirming its stability. This binding pose was mediated by the same residues detected to be key for sliding in living organisms (Lys3, Lys18, and Lys83) [25], which indicates that this is the search state. On the other hand, the binding with the arms and saddle represents the recognition state. Hence, our simulations offer a structural model of the three diffusion rates detected experimentally [29], where the slowest mode would involve sliding with the recognition state, the intermediate mode sliding with the searching state, and the fastest mode would involve hopping via the *β*-ribbon arms (Figure 6).

In addition, our simulations captured the transition from the non-specific to the specific state only in the presence of damaged DNA. Its enhanced flexibility relative to B-DNA facilitated this transition, making any barriers practically negligible. We discovered a “concerted” mechanism between the “induced fit” and “conformation selection” scenarios, in which the action of HU is required to achieve the strong DNA bending of the final complex, but DNA deformability is required for the protein to identify its target. These results are consistent with a previous experimental study that showed that the association between a protein and DNA is not only limited by diffusion, but also by their ability to form the high-affinity complex once they found one another [1].

Notably, we did not see direct binding to the specific-binding region, which is the most positively charged, due to its inaccessibility behind the flexible *β*-ribbon arms (see Figure 3B). Instead, our simulations revealed that HU attaches to DNA in a sequence of steps, progressing from the encounter complexes to the high affinity complex via a mechanism of the protein rolling or reorienting over DNA. The benefit of making the most positively charged region inaccessible could be to prevent the protein from becoming trapped at any DNA location. Instead, this multistep mechanism of binding, along with the requirement of DNA bending, would guarantee that the high-affinity complex only forms at the DNA target location.

We anticipate that the same principles discovered here will apply to other DNA-binding proteins, especially architectural ones, characterized by their substantial positive charge and capacity to distort DNA. The electrostatic interactions between these proteins and DNA would be precisely modulated to optimize both their search kinetics and the stability of the recognition state. The presence of disordered regions adjacent to the most positively charged areas, as seen in the yeast protein Nhp6A and human p53 [11, 29], may represent a shared approach to enhance their inaccessibility and prevent entrapment. Moreover, these interactions may be associated with the DNA’s flexibility, potentially serving as a recognition signal, thus accelerating target identification.

## Author contributions statement

E.W.C., M.C.L. and A.N. conceived the simulations; E.W.C. conducted the simulations; E.W.C. and A.N. analysed the results; E.W.C., M.C.L and A.N. wrote and reviewed the manuscript.

## Acknowledgements

We would like to thank J. A. L. Howard for useful discussions. This work was supported by WW Smith Fund (E. W. C.); EPSRC EP/N027639/1 and EP/Y008693/2 (A. N.); as well as EP/T022205/1, EP/X035603/1, EP/P020259/1, and EP/T022167/1 for computational resources. We thank the University of York for the Viking cluster, the local high performance computing facility, and for providing pump-priming resources. Simulations are available at University of York Data Repository (DOI:10.15124/033aa782-750e-4d19-9909-cdb2ff86ddc9)

